# Community ecology of phages on a clonal bacterial host

**DOI:** 10.1101/2023.09.01.555216

**Authors:** Nora C. Pyenson, Asher Leeks, Odera Nweke, Joshua E. Goldford, Paul E. Turner, Kevin R. Foster, Alvaro Sanchez

## Abstract

Bacteriophages are the most abundant and diverse biological entities on Earth, yet the ecological mechanisms that sustain this extraordinary diversity remain unclear. Here, we have discovered a general mechanism that allows phage diversity to outstrip the diversity of their bacterial hosts. We assembled and passaged dozens of diverse phage communities on a single, non-coevolving strain of *Escherichia coli* until the phage communities reached equilibrium. In all cases, we found multiple phage species coexisted stably, despite competition for a single, clonal host population. Coexistence within these communities was supported through host phenotypic heterogeneity, whereby phages specialized on cells adopting different growth phenotypes. Our experiments reveal a rich community ecology of bacteriophages where multiple phage species coexist and interact at the scale of a single bacterial host.

**One-Sentence Summary:** Diverse communities of phages emerge stably and consistently on a clonal bacterial host, enabled by phenotypic heterogeneity.

## Main Text

Bacteriophages are central to the functioning of many microbial communities (*1, 2*), from global carbon cycling in marine ecosystems (*3*) to the health of the human microbiome (*4*). These ecological functions are underpinned by the extraordinary abundance and diversity of bacteriophages. There are an estimated 10^31^ phage virions on Earth (*5*), with entirely novel taxonomic groups of phages frequently uncovered through metagenomics (*6, 7*) and electron microscopy (*8*). It is typically assumed that this phage diversity has arisen through host genotypic diversity, with each phage specializing on a genetically different host strain (*9*). A null ecological expectation, grounded in the competitive exclusion principle, is that each clonal host population could only maintain a single phage, as each host type represents a single resource whose consumers are competing phage species (*10, 11*). This expectation would predict that the coexistence of multiple phage species in the same environment requires a genetically diverse host population. Consistent with this logic, experimental work using multi-species bacterial communities (*12, 13*) or co-evolved host species (*14, 15*) has shown that host genotypic diversity supports phage coexistence.

However, phage taxonomic diversity substantially outstrips the diversity of their bacterial hosts across a wide range of habitats including the human gut (*16, 17*) and oral cavity (*18*), the ocean surface (*19, 20*), and hot springs (*21*). For example, the infant gut contains over two thousand DNA phage species and around two hundred bacterial species (*22*). Phage diversity can even be seen at the single-cell level; in some environments, the majority of infected bacterial cells contain two or more distinct phage species (*23, 24*). In these environments, phage diversity cannot be explained by host genetic diversity, suggesting that other mechanisms must be at play. Despite the central role that phages play in microbial ecology, we still understand surprisingly little about the processes that drive and maintain this diversity (*25, 26*) and we lack direct empirical tests of whether host genetic diversity is indeed a requirement for multiple phage species to coexist (*25, 27*). To resolve this fundamental question, we investigated whether diverse populations of naturally-isolated phages could coexist when passaged on a single non-coevolving bacterial host of *E. coli* under stable and well-controlled laboratory conditions.

## RESULTS

### Lytic phages assemble into diverse communities

To test whether competitive exclusion generically limits phage diversity to a single phage species per host strain, we first isolated a diverse collection of naturally occurring phages capable of infecting the *E. coli* K-12 strain BW25113 (Fig. 1A) before performing top-down community assembly experiments. We isolated *E. coli* phages from different environmental samples, like animal droppings and river water (collected in New Haven, Connecticut, USA) that generated clearings on a lawn of *E. coli*. Individual phages were then plaque purified and their genomes were sequenced. This process resulted in a collection of 27 double-stranded DNA (dsDNA) tailed bacteriophages (Caudoviricetes) (*28*) that contained substantial genetic, life history, and taxonomic diversity. Most phages had distinct plaque morphologies (Fig. 1B) and shared very limited homology with one another, apart from phages in the same family (Fig. S1). This genetic divergence reflected the fact that our phages were taxonomically diverse (*29*) (Fig. 1C, Table S1), containing 10 families, 13 genera, 24 species, and 27 strains (*30*). Our collection contained 14 obligately lytic (aka lytic) phages, meaning that they lyse infected host cells, and 13 temperate phages, meaning that they can integrate into the bacterial genome (Fig. 1C, Fig. S2).

**Fig. 1.**
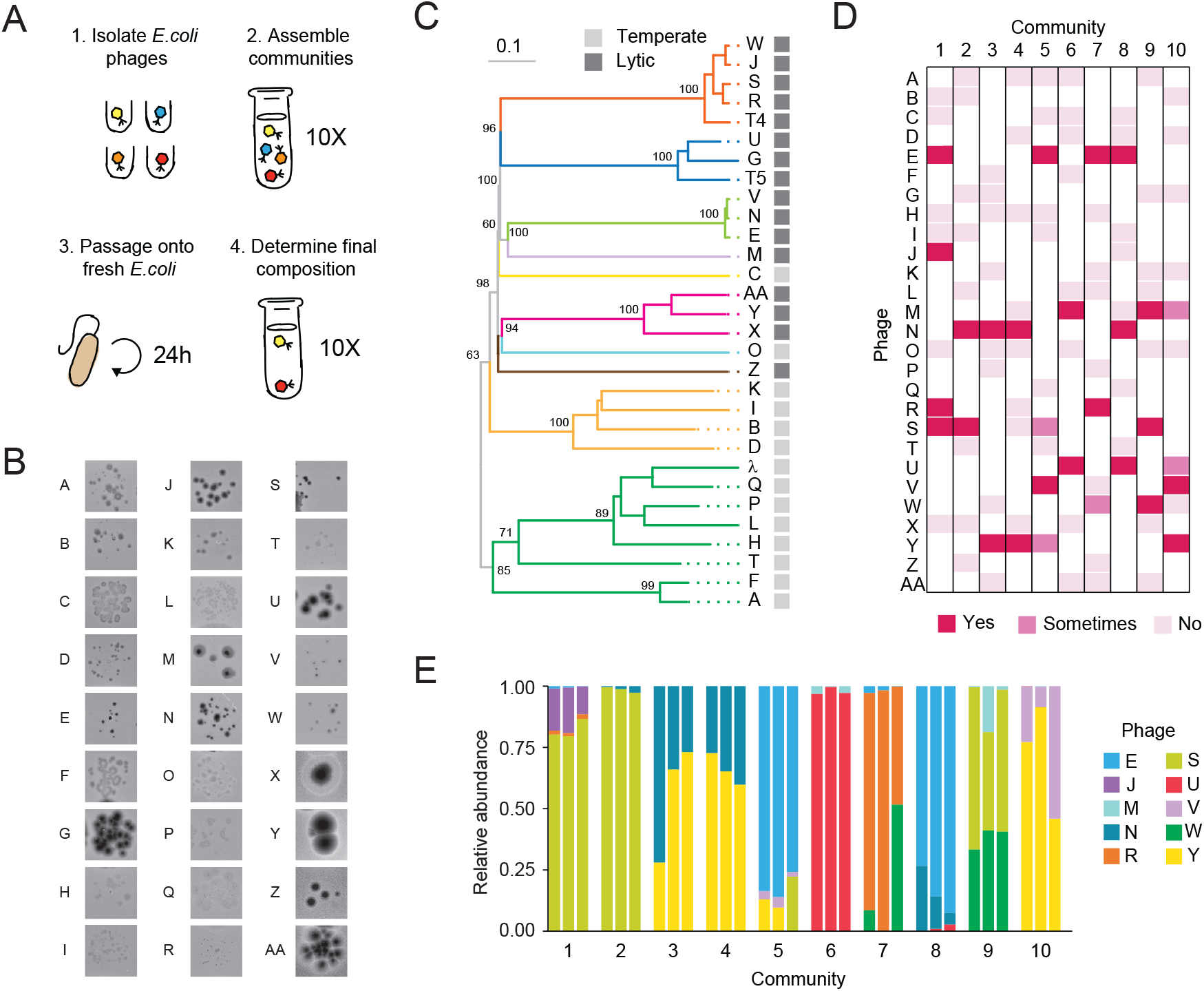
Phage communities are diverse, common, and predictable on a single host. (**A**) Illustration of the experimental workflow. 1) Isolate and genome sequence 27 *E. coli* phages from natural samples. 2) Infect *E. coli* with 10 different starting communities each containing a random set of 10 phages from our collection. 3) Filter, dilute, and passage the phage community onto fresh bacteria every 24 hours. 4) At day 12 determine the composition through deep sequencing and plaque morphologies. (**B**) Diverse plaque morphologies of the 27 phages in our collection. (**C**) Phylogenetic tree including all 27 phages in our collection and model phages T4, T5 and lambda as references. The numbers above the branches are GBDP pseudo-bootstrap support values from 100 replications and the branch length is scaled by the D0 formula. The branches were colored based on family classification (see Fig. S1 or Table 1 for family names). (**D**) Every community contained multiple diverse phage strains at the final passage, as determined through deep sequencing. “No” indicated that the phage was present in the starting community but not detectable at the final passage in any replicate. “Sometimes” indicated that the phage was present in some replicates, while “Yes” indicated that the phage was present in all replicates. (**E**) The abundance and identity of phage strains found through deep sequencing varied among different communities, but was similar for the three replicates of the same community.

With this diverse collection of phages we assembled 10 different phage communities, each containing a random subset of 10 different phage strains from the collection, and then tested whether they could coexist on a single non-evolving host population (Fig. 1A). Three replicates of each phage community were passaged onto fresh *E. coli* every 24 hours and the phage component was filtered before each passage to eliminate any surviving bacterial cells. At the 12^th^ passage we determined which phages remained by shotgun sequencing the final community, assembling phage genomes de novo, and then comparing the assembled genomes to our collection of 27 phage species (Methods). We validated that the deep sequencing accurately measured the relative abundance and identity of phages by quantifying plaque morphologies through top agar plating of three communities (RMSD = 0.17, N = 9) (Fig. S3).

We found that all 10 final phage communities maintained ecological diversity, with two to four coexisting phage strains from at least two different families (Fig. 1D, E, Fig. S4A) in every community. Replicate communities started from the same initial phage community resulted in highly similar final communities. Only three of the ten inoculated communities produced replicates that differed in composition (communities 5, 7, and 10), and these communities only differed in one out of their three coexisting phages. Communities that were started from different initial phage communities produced final communities with different sets of coexisting phages (a high beta diversity), with just two exceptions (communities 3 and 4). We found that the emergent diversity depended on phage life history: all remaining phages were lytic, while temperate phages were never present in the final passage (Fig. S4A). This diversity was found despite various features of our experimental set-up that were intended to limit the potential for phage coexistence: i) a single, clonal strain of *E. coli* was the only host; ii) the host population was removed at each passage so it could not co-evolve and diversify throughout the experiment; and iii) phage communities were diluted at the beginning of each passage, which should disfavor phage species that did not replicate or that were not competitive.

To test whether our finding of generic multi-phage coexistence on a single host was robust, we repeated the experiment with the following modifications to our protocol: (i) allowing for host co-evolution; (ii) reducing the starting phage diversity. When the same experiment was performed without filtering the samples between each passage, which would allow for spontaneously resistant mutant cells to carry over between passages, we again found phage coexistence in all 6 tested communities (Fig. S4B). Similarly, diverse phage communities were ubiquitous when we started each passaging experiment with only 5 phage strains instead of 10 (Fig. S4C). In total, we found that 13 out of 14 of the lytic phages from our collection were present in at least one community from these and our original experiments (Fig. S4, Table S1). This suggested that virtually all of our lytic phages had the potential for coexistence, albeit not in every community. However, none of these communities contained any temperate phages, which additional experiments suggested was caused by competition from lytic phages: temperate phages were able to survive the passaging regime in monoculture at high titers above the limit of detection for our communities (Fig. S5A, B).

### Coexisting phage communities emerge rapidly, are robust to perturbations, and are dominated by negative interactions

To investigate how quickly our communities assembled throughout the passaging regime we quantified phage abundances using plaque morphologies for four representative communities from Fig. 1 starting at the first passage. The most abundant phages at the final passage usually emerged as the dominant members of the community by passage 3 and remained dominant until the final passage (Fig. 2A). In addition, most phages that were absent from the final communities were below the limit of detection by passage 3. These results are consistent with rapid ecological dynamics that arose early during passaging and likely reflects significant differences in growth rates between phage species. However, in a few cases, including the replicates of community 7, the phages in the final community did not emerge until later passages, revealing longer timescale processes. In all communities, the absolute population size of the phages tended to fluctuate, but the overall diversity and species composition remained relatively similar from early in the experiment.

**Fig. 2.**
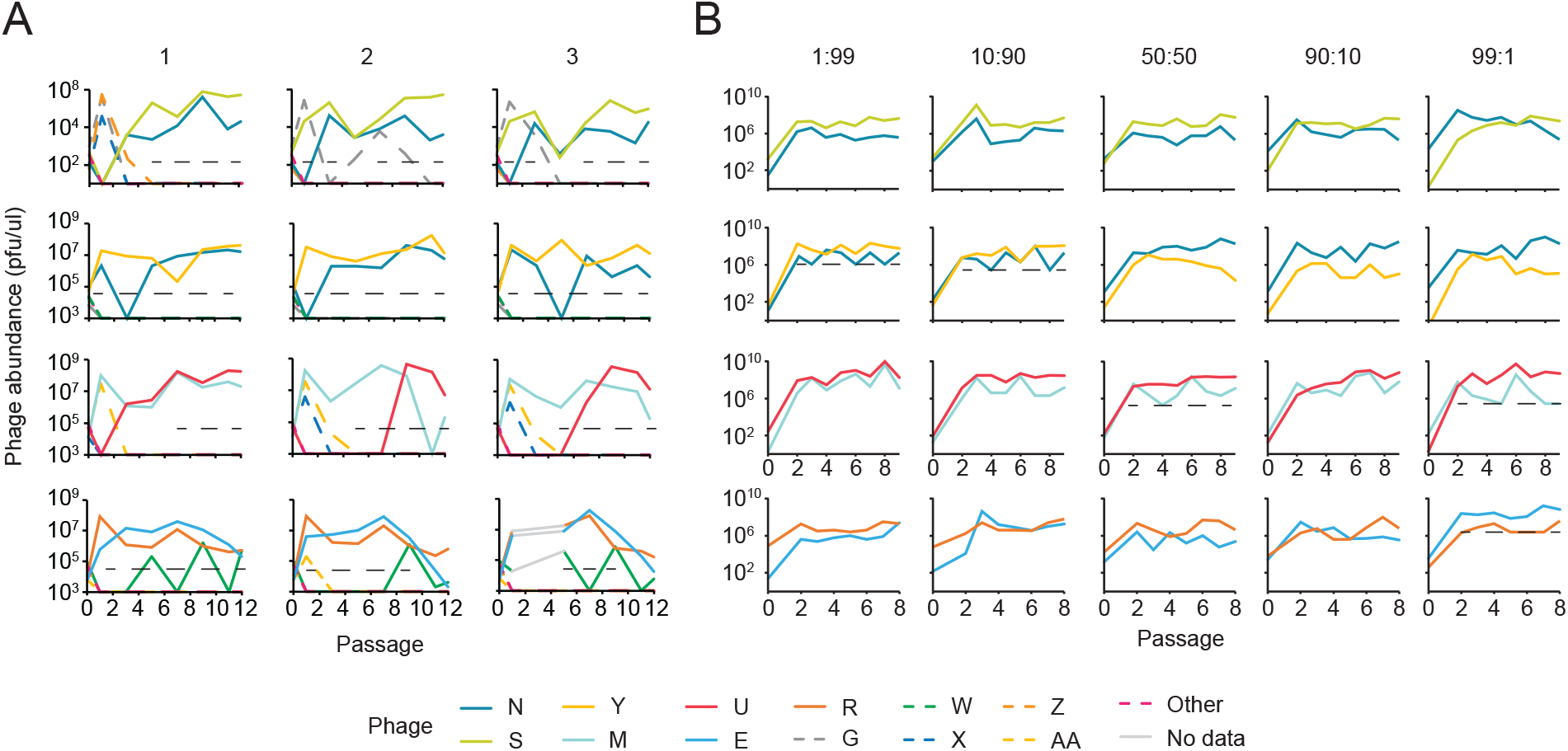
Diversity is stable throughout community passaging. Phage abundances over time for communities 2, 3, 6, and 7 (top to bottom), quantified using plaque morphologies. (**A**) The three replicates (left to right) of four communities from Fig. 1D show the early dominance of final community members. The average limit of detection for each replicate is indicated by the black dotted line or is otherwise indicated by the x-axis. Phages that are represented with a dashed line were absent from the final community. Phages were quantified on passage 0 and then every other passage starting at passage 1 until passage 12. (**B**) Reconstituted 2-member communities maintained both phage species irrespective of starting abundance. The initial abundances of each phage strain (listed from left to right) varied from ∼1% to ∼99% of the total phage population for community 2 (phage N: S), 3 (N:Y), 6 (M:U), and 7 (E:R). For simplicity we reconstituted community 7 with only phages E and R, although the original communities also had phage W. The limit of detection is only shown for replicates where one of the two members was undetectable at least once during the passaging. Phages were titered on passage 0 and then every day starting at passage 2 until passage 9 (communities 2, 3, 6) or passage 8 (community 7).

The speed at which our community composition became fixed suggested that our phage communities would be stable to perturbations. We directly evaluated the stability of our communities by testing whether each phage could invade the community when initially present as a rare proportion of the population. We reconstituted and passaged the four different 2-member communities examined above (Fig. 2A) and varied the starting proportion of each phage between 1%, and 99% of the total population (Fig. 2B). We found that both phages were still detectable at the last passage in every condition for all 4 communities, with the exception of one replicate of community 6. All phages gained an increased relative growth rate when starting from rarity (negative frequency dependence; Fig. S6), that was sufficient to maintain coexistence following perturbations in the relative abundance of each species. These results demonstrated that our phages displayed the hallmarks of stable coexistence (*31*).

Our results show that the same communities of phages emerge rapidly, predictably, and stably. The properties and functioning of a stable community can be understood through species interactions (*32*): ie. how one species affects the growth and reproduction of other species (Fig. 3A). We determined the direction of interactions for a range of 2-phage (Fig. 3B) and 3-phage (Fig. 3C, D) communities by comparing each phage’s abundances when grown in a community compared to when grown alone. This experiment revealed that pairwise interactions were typically negative with examples of both competition (-/-) and ammensalism (0/-) among the phages. None of our phage pairs showed signs of mutualism, but one interaction was asymmetric: phage N benefited from co-culture with phage S, while phage S did worse in co-culture (+/-) suggesting an exploitative interaction. The predominance of negative interactions in our communities is consistent with the rapid exclusion of many species early in our passaging and with the finding that our communities are robust to perturbation, as negative interactions can stabilize coexistence (*32*).

**Fig. 3.**
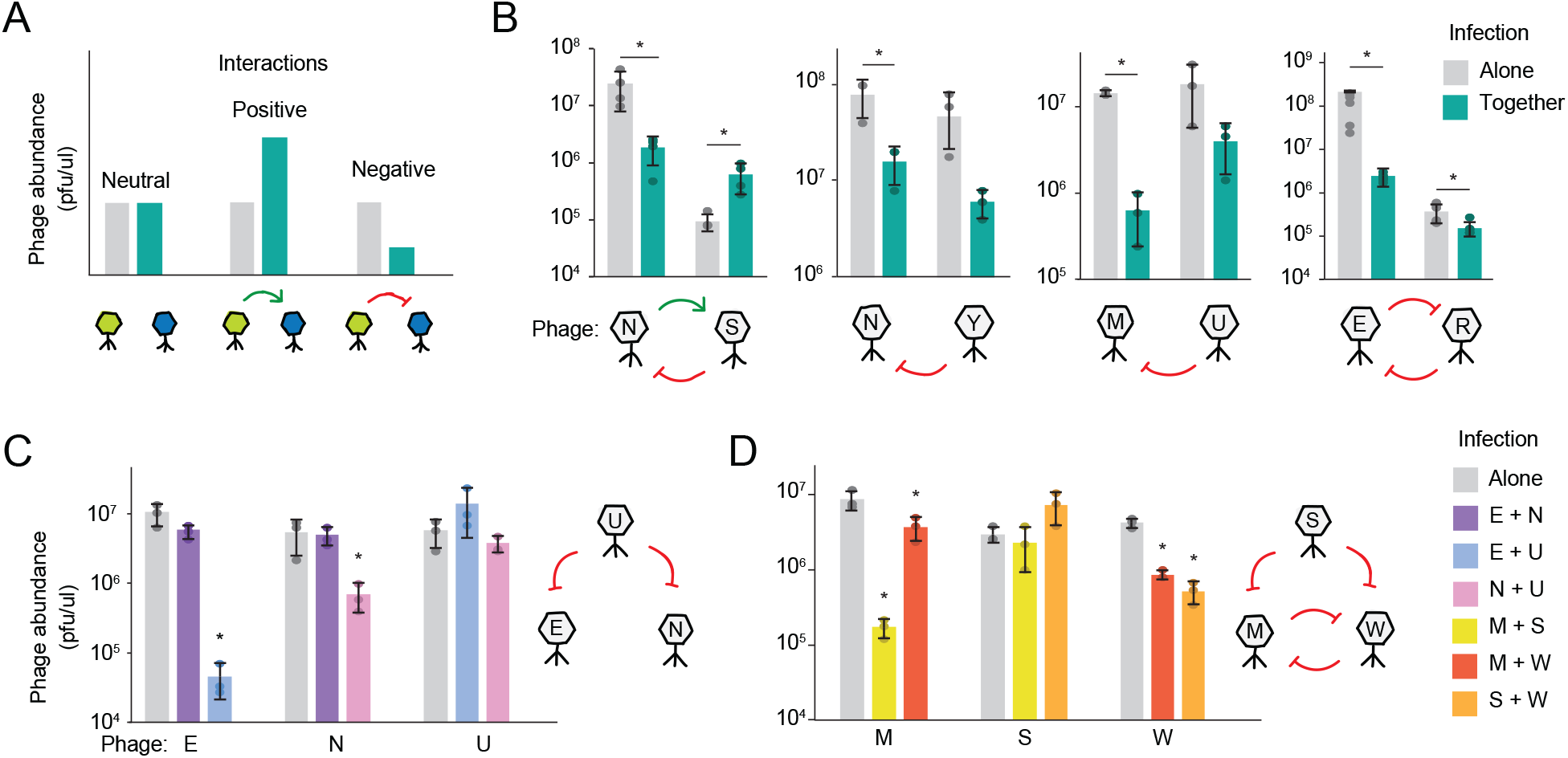
Phage-phage interactions are largely negative. (**A**) Example results that show how social interactions are determined through changes in phage abundance. The green phage can either have no impact, increase, or decrease the blue phage’s abundance during coinfection, relative to when the blue phage is infecting alone. These interactions would be neutral, positive, or negative, respectively. All samples were passaged for 3 days to allow populations to reach equilibrium before measuring the phage abundances through top agar plating. (**B**) Phage interactions for 2-phage communities 2, 3, 6 and 7 (left to right) were determined by comparing a phage’s abundance during an infection together (2-phage) versus alone (1-phage). Results are the mean of 3 to 6 biological replicates ± SD. Starred samples had *P* values under 0.05 (from left to right: 0.03, 0.02, 0.03, <10^-4^, 0.01, 0.01). Under each community is a graphical illustration of the social interactions with green arrow arcs for positive interactions and red inhibition arcs for negative interactions. (**C** and **D**) Same as (B), but showing pairwise interactions within 3-member (C) community 8 or (D) community 9, instead of 2-member communities. Starred samples had *P* values under .05 (comparisons against culturing alone, from left to right: 0.008, 0.04, 0.004, 0.03, <10^-3^, <10^-3^). All *P* values were calculated with a two-tailed T test.

### Productive coinfection of individual host cells is rare

One way that stable coexistence can emerge between competing species is via niche partitioning of their shared resource. Coexisting phage species outnumbered the bacterial host cells (ie. had a high multiplicity of infection, or MOI) at the onset of virtually every passage during our community experiments (Fig. S7), which suggested that multiple phages could be sharing resources and niche partitioning within the same coinfected cells. To test whether this high MOI resulted in productive coinfections, we used single-cell flow cytometry sorting to study the diversity of phages produced by single bacterial cells after infection (*33, 34*) (Fig. S8, Methods). For two different MOIs (3.9 and 4.8) we found that the proportion of cells producing both phages from community 2 was negligible (<10%), and far lower than the expected proportion of coinfected cells (92 -100%, based on the joint Poisson distributions given the MOI for each phage). Productive coinfections were also negligible for cells infected with a different set of coexisting phages at MOIs of 25 and 14. These experiments revealed that it was rare for multiple phage species to emerge from the same cell, showing that phage coexistence was not achieved by processes that require successful coinfection of individual cells.

### Phenotypic heterogeneity within the host population supports phage coexistence

Given that each phage species was successfully infecting different cells within the host population (Fig. S8), we next explored whether coexistence could emerge through niche separation between different host cells. The bacterial hosts in our experiments were isogenic, but cells can differ in their physiological states through phenotypic heterogeneity (*35*). One well-documented form of phenotypic heterogeneity among our normal host population (a 24-hour old *E. coli* culture diluted into fresh media) is the growth variability between different cells (*36, 37*). The coexistence of multiple phage species could be supported by phenotypic heterogeneity if such cell-cell differences resulted in differential phage infection. This process would be akin to viral coexistence in multicellular hosts where each viral species specializes on a different tissue or cell type (*38, 39*). We focused on community 2 for the remaining experiments since these phages showed a particularly consistent composition over time (Fig. 2B).

We first tested whether changes to host growth potential of the entire culture affected relative phage growth rates. To do this we infected a 3-hour, 24-hour, or a 72-hour culture of *E. coli* since increasing or decreasing the culture time causes slower or faster growth, respectively, upon dilution into fresh media (*38, 39*). We found that changing the culture age completely reversed which phage species was dominant in the community. As the age of the culture increased from 3 to 24 to 72 hours, slowing the bacterial growth, the community composition flipped from an average of 95% to 66% to 15% phage N (Fig. 4A). These results, generated after coinfection of each culture, were consistent with the productivity of each phage when only a single species was present (Fig. 4B) and suggested that fast growing bacteria preferentially produced phage N and slow growing bacteria produced phage S.

**Fig. 4.**
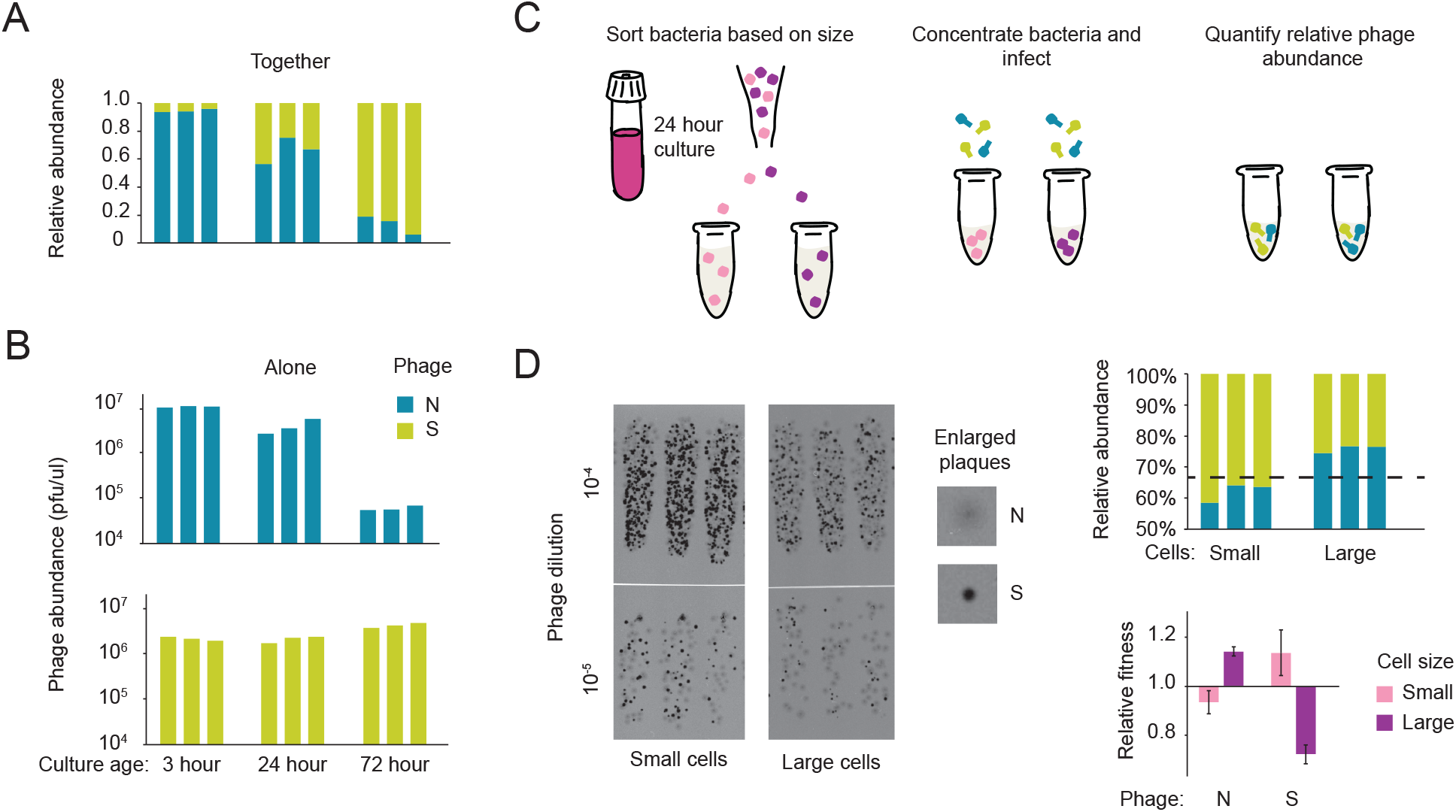
Host heterogeneity supports phage diversity. (**A**) Phage relative abundances are controlled by host physiology. Cultures of different ages were infected with community 2 after passaging the community for 2 days to allow the phage composition to equilibrate. Phage abundance was measuring through plaqu-ing. Results shown are biological replicates. The *P* values were less than 0.05 for 3 hour vs 24 hour (0.006) and 72 hour vs 24 hour (0.001). (**B**) Same as (C) except cells were infected with either phage N or phage S, instead of both phages, and absolute abundances are shown. The *P* values were less than 0.05 for phage N (top): 3 hour vs 24 hour (0.001), 72 hour vs 24 hour (0.01) and phage S (bottom): 72 hour vs 24 hour (0.004) and 72 hour vs 3 hour (0.003). (**C**) Schematic for determining the phages produced by small and large cells: cells from a 24 hour culture were separated by size, reconcentrated and infected with phage, then the abundance of phages was determined after incubation. (**D**) When coin-fected with community 2, small and large cells preferentially produce phage S or N, respectively. Top agar plating of phages produced by cells of different sizes (left panel), relative abundance of each phage, with the dotted line indicating the starting proportion of phage N (right panel, top), and the fitness of each phage by cell size, relative to that phage’s starting proportion (right panel, bottom). Results are the mean of three biological replicates ± SD with *P* = 0.002. See Fig. S10 for an additional replicate experiment. All *P* values were calculated with a two-tailed T test.

We next used flow cytometry methods to validate that the difference at the culture level were reflected in phage specialization on different growth phenotypes within a single culture. We separated the slow- and fast-growing subpopulations from the 24-hour culture by using relative cellular size (forward scatter), since shrunken (small sized) *E. coli* cells from an overnight culture grow slower than normal (large sized) cells (*39*–*41*) (Methods). After confirming that small cell size was a marker for slow growth (Fig. S9), we sorted and infected 20K small and large sized cells with phages from community 2 (Fig. 4C, Methods). This experiment showed that each phage species performed better on a different cell type (Fig. 4D), in the direction that is consistent with our previous culture-level findings (Fig. 4A, B). Phage N, which did better on the 3-hour culture (fast growth) also performed 15% better on large cells (fast growth) (95% CI [12.9, 17.2]), while phage S, which did better in the 72-hour culture (slow growth), performed 11% better on smaller cells (slow growth) (95% CI [1.1, 21.7]), relative to their starting population. We found the same effect in a replicate experiment, with phage N performing 16% better on large cells (95% CI [10.4, 23.5]) and phage S performing 10% better on small cells (95% CI [-6.23, 27.7]) (Fig. S10). These results show that even within a clonal host population, there is sufficient phenotypic heterogeneity to allow niche separation of phages onto different cellular sub-populations, allowing coexistence between taxonomically diverse phage species.

## DISCUSSION

Despite the importance of phages in a wide range of Earth’s microbiomes (*1, 3*), we still understand little about the ecological mechanisms that support phage diversity (*25, 27*). By infecting a clonal *E. coli* population with diverse communities of phages, we have discovered that multiple lytic phage species generically coexist together on a single host. This coexistence was ubiquitous and taxonomically diverse; not a single one of our communities was reduced to just one phage species, or even one family of phages, even after 12 passages. We show that these communities are ecologically stable, whereby phages can establish and persist from a rare proportion of the population. Within these phage communities, as in many other microbial systems (*32, 42*), ecological interactions were dominated by competition and amensalism. Moreover, we find evidence of niche separation within the communities that enables coexistence in spite of competition. Each phage species could persist in the community by differentially replicating on host cells with different growth phenotypes (ie. slow or fast growth) that spontaneously arose within the *E. coli* culture.

Our work provides a new null expectation where phage diversity is not generically constrained by the number of host genotypes. This contrasts with the typical assumption that the broad phylogenetic diversity of phages found in nature is supported by the genetic diversification of their bacterial hosts (*9, 25*). This expectation is rooted in consumer-resource models (*43, 44*) where the diversity of coexisting consumers (ie. phages) is capped by the diversity of available resources (*45*). Our work is consistent with studies in other microbial systems that have found that the nature of resources is often more subtle than it may appear: given enough starting diversity, multi-species coexistence arises in bacterial communities, even when cultured on a single growth-limiting resource (*46*). From the perspective of the phages, our ubiquitous findings of multi-species communities suggests that both ecological and evolutionary dynamics, like recombination and host specialization, will be shaped by diverse phage coexistence (*32, 47*–*49*).

We found that variations in host physiology were critically important for our phage communities, dictating the relative abundance of different phage species and supporting phage coexistence on an isogenic host population. Even among clonal, laboratory-grown *E. coli* populations, phenotypic heterogeneity arises spontaneously through molecular fluctuations in transcription and translation and is therefore unavoidable (*35*). Natural systems are expected to have higher levels of cell-cell heterogeneity than laboratory culturing (*50*), and so we expect that the levels of phage diversity supported by a single bacterial genotype is likely even higher in nature than what we find in this study. These findings connect phage ecology with the ecological theory of mammalian virus coexistence (*38, 39*), where multiple viruses can persist on a single host by specializing on different cell types or tissues.

Just as bacterial communities, like the gut microbiome, can form at the scale of a single, multicellular host, we show here how phage communities can form at the scale of a single bacterial strain. The ecological frameworks used to understand bacterial communities (*32*) served as our guide for characterizing how factors like life history traits, inter-species interactions, and resource partitioning shape our phage communities. These results reveal how, even under highly constrained laboratory conditions, competitive exclusion does not generically limit the diversity within complex phage communities.

## Acknowledgments

We would like to thank members of the Sanchez, Turner, and Foster Labs for their helpful discussion.

## Author contributions

Conceptualization: NCP, KF, AS; Methodology: NCP, JEG, AL, AS; Formal Analysis: NCP, AL; Investigation: NCP, ON; Writing – original draft: NCP, AL; Writing – review & editing: NCP, AL, PT, KF, AS

## Data and materials availability

All the data from this study are available from the authors upon request.

## Supplementary Materials

### Methods and Materials

Figs. S1 to S12

Tables S1

References (*51*–*62*)

