## Supplementary for "Community ecology of phages on a clonal bacterial host"

**The PDF file includes:**

Materials and Methods
Figs. S1 to S12
Tables S1
References (*51-62*)

### Materials and Methods

Bacterial culturing and infection

*E. coli* strain BW25113 was used for all experiments with incubations at 37°C. Culturing of uninfected bacteria was done in 3ml of LB broth (Miller) (American Bio) in a 15ml snap-cap tube (Falcon) grown with shaking. Frozen stocks of *E. coli* were made with 40% glycerol (final concentration) and kept at -80°C.

Infections were carried out in 300µl LB + salts (final concentration 100mM CaCl2 and 100mM MgCl2) in either 2ml 96-deep-well plates (Celltreat) sealed with piercible aluminum seal (Thomas Scientific) or in 1.5ml screw cap tubes. For 96-well plate infections, we spatially isolated each sample by leaving 2 empty wells in all directions to avoid cross contamination between communities. We used a fixed concentration of each phage for the first passage of an experiment (Day 0), starting with a total of 100K pfu with equal proportions of all phages (unless otherwise noted for some 2-phage communities). For 10-phage communities, the members were selected randomly, while for communities consisting of 5 different phages, the starting communities were selected semi-randomly to include at least 2 lytic phages.

Phage was added to the sample immediately after adding a fixed volume or concentration of 24-hour old *E. coli* culture (more on this below). The infections were incubated for 24 hours then the phage was collected by filtering 100µl through a 96-deep-well 0.22 µm filter plate (Acroprep) and the unused portion of the sample was stored at -80°C. We added 10µl of this filtered phage (dilution of 1:30) for infection of the next passage. For community assembly experiments (Fig. 1, Fig. S4), passaging was continued until day 12 while for community stability tests (Fig. 2B), passaging was continued to day 9. For most other assays that required infecting cells with a phage community (ie: phage interactions (Fig. 3), infection growth curves (Fig. S11), and niche partitioning (Fig. 4)) we performed experiments using community samples that were passaged for 2-3 days to allow the composition to stabilize.

We used two different techniques for inoculating the fresh "overnight" 24-hour culture. For community assembly experiments and stability tests, *E. coli* was streaked onto a fresh LB agar plate before the first passage and a single colony of *E. coli* was used as inoculum for the 24-hour culture that was started before each passage. This plate was stored at 4°C and a single colony was used as inoculum for the remainder of the experiment. At the onset of each passage the 24-hour culture was diluted 1:100 and immediately infected with phage. To increase the consistency of phage titers (*51*, *52*) throughout our study we slightly modified the inoculation and infection protocol used in all other experiments. Instead of starting a 24-hour culture with a single colony, the culture was inoculated by scraping the frozen *E. coli* stock. We also used a fixed number of cells at each passage instead of a fixed volume for all other experiments. We measured the OD600 before each infection to calculate the bacterial concentration where an OD of 1.0 is 8 x $10^6$ colony forming units (cfu)/µl. We then added bacteria at 4 x $10^4$ cfu/µl (final concentration). In general we found that a 24-hour *E. coli* culture had an OD between 4.5 and 6.5. The previous technique (diluting bacteria 1:100) would have given the same concentration of bacteria from an OD of 5. Using the frozen bacteria as inoculum gave community 2 more stability (Fig. S12) than when passaged on bacteria started from a colony inoculum (Fig. 2).

Bacteriophage isolation

Phages were isolated from different environmental samples collected between September and October 2020 from New Haven (Connecticut, USA). Each sample was collected in a sterile 1.5ml tube and briefly vortexed. To increase the abundance of *E. coli* phages, 50µl of an overnight culture and 500µl of LB + salts were mixed with each sample (*53*). After an overnight incubation without shaking the supernatant from each sample was then filtered to remove the bacterial component. To check for the presence or absence of phages in the sample, we plated 2µl of the sample on a top agar plate. Samples that generated a transparent spot on the lawn of *E. coli* were then used for two rounds of single plaque purification. Single plaques from the last round of purification were then used to make stocks of each phage that were subjected to genome sequencing. Out of 172 environmental samples collected we were able to single plaque purify phages from 77 samples. We subjected 48 phages to sequencing and chose 27 phages for inclusion in our collection.

Bacteriophage handling and culturing

High-titer stocks of each phage were generated by mixing a single plaque with a 24 hour culture that was diluted 1:20 into 600µl LB + salts. After 24-hour incubation, phage was passed through a 0.22 µm filter to remove any surviving bacteria. Working stocks of each phage were stored at 4°C and discarded after 2 weeks. Permanent phage stocks were frozen with 40% glycerol (final concentration) and stored at -80°C.

Top agar preparation and plaque quantification

For plaque quantification we used the double-agar overlay method. The bottom layer was comprised of LB plates made with molecular-biology grade agarose at a final concentration of 0.8% w/v. We found that higher quality agarose, as compared to agar, gave a smoother surface for plating phages when used in the bottom and top layer of the plate. The LB + salts top agar (final concentration 100mM $CaCl_2$, 100mM $MgCl_2$, 0.4% w/v agarose, 5% glycerol) was stored at 55°C and contained glycerol to improve plaque visualization (*54*). To quantify plaques we mixed 6ml of top agar and 150µl of a 24-hour culture and poured onto the surface of the LB agarose plate. Serial dilutions of phages were done in 1X PBS and then plated by spotting or dripping a small volume onto the surface of the top agar. We typically plated 5µl-10µl of 10-fold dilutions from $10^{-3}$ to $10^{-8}$. After overnight incubation, plates were scanned using the Epson Perfection V850 Pro Scanner at 800dpi using the transparency setting. Scanned images were then imported into Adobe Illustrator to quantify plaques and visually distinguish plaque morphologies of different phages.

For some experiments we changed the top agar recipe to enhance plaque visualization. To reduce the plaque size of phage Y we increased the concentration of agarose (0.8% w/v). To increase the size of phage R plaques we omitted the glycerol and doubled the volume of overnight culture used (300 µl) but did not change the agarose concentration.

Bacteriophage DNA isolation and sequencing

Genomic DNA from 100µl of phage stocks or 50µl of phage communities was purified using the ZR-96 Clean and Concentrator Kit (Zymo) according to the manufacturer's instructions. We excluded the dnase or protease treatments that are sometimes suggested for phage DNA extraction (*53*, *55*). Each sample was prepped and sequenced at Seq Center using their Illumina NextSeq platform.

Isolated phage genome assembly

To generate full genomes from isolated phage genomes we first trimmed the fasta files using Sickle (https://github.com/najoshi/sickle). Full contigs were then assembled using SPAdes (*56*) with default parameters, but using only the first 400,000 lines of the fasta files, since excessive coverage leads to problems with assembly (*57*). We verified the assembly of a single contig through Bandage and got ~40 fold coverage per phage genome.

Pairwise genome comparisons and lifestyle characterization

Pairwise genome comparisons were performed using the default settings on Virdic (*58*) on the web application (http://rhea.icbm.uni-oldenburg.de/VIRIDIC/). Phages were assigned as lytic or temperate traits by combining analyses done both bioinformatically and experimentally. For bioinformatic analysis we used two tools, submitting genomes to PhaBOX (the PhaTYP (*59*) module) (https://phage.ee.cityu.edu.hk/) web application and with the classifier BACPHLIP (*60*). The predictions from these tools (Table S1) agreed with experimental assays looking at spot turbidity and growth curves of infected cells (Fig. S2). We determined spot turbidity by plating a concentrated phage stock on a lawn of *E. coli*.

Phage community sequencing analysis

To identify phages from a community we first assembled the sequencing reads into individual contigs as described for isolated phage genomes, except we used the whole fasta file for assembly to ensure coverage of rarer phage strains. We identified phages within the community by using a local NCBI blastn search (word size of 1000) to compare each community contig against a local database with our collection of 27 phage genomes. We used the average coverage of each contig in the community to calculate the relative abundance of each phage in the community. None of our phage communities showed evidence of cross contamination from other communities. For the most abundant phage in a community, this process gave 1000 – 4000 fold coverage of the entire genome. Our limit of detection for a phage through deep sequencing was ~0.1% of the total phage population or ~1X coverage of a phage's genome.

In communities with multiple *Straboviridae* phages we used a modified NCBI blastn search to identify the presence of each phage from this family since these phages had high homology with one another. In communities with multiple *Straboviridae,* only the most abundant *Straboviridae* phage genome was assembled into a single contig. The lower abundance members were assembled into many smaller contigs (100-1000 base pairs) containing only the regions of non-homology with the more abundant *Straboviridae*. To identify phages from the smaller contigs we made a local database containing a unique sequence from each phage in the *Straboviridae* family. We then aligned our community contigs to this database using NCBI blastn (word size of 100). We used this same process to identify multiple *Stephanstirmvirinae* phages from the same community.

Taxonomy and phylogenetic tree

Each phage was given a shorthand name used in the paper as well as a formal name (Table S1) according to the rules of the International Committee on the Taxonomy of Viruses (ICTV) (*28*). The phylogenetic tree was generated using VICTOR (https://ggdc.dsmz.de/victor.php) (*30*) with the formula D0 producing an average support of 65%.

To identify the taxonomic classification of each phage we first performed whole-genome blastn searches against the virus database (https://blast.ncbi.nlm.nih.gov/Blast.cgi) to find classified phages with high identity. Additional taxonomies were identified using PhaGCN (https://phage.ee.cityu.edu.hk/) (*61*) and the aforementioned VICTOR analysis. To classify phages in the same genus or species we used a cut-off of 70% or 95% nucleotide identity across the full genome length (*62*).

Infection growth curves

Bacterial growth curves were performed in triplicate using 96-well plates with 150µl LB + salts in each well with 4 x $10^4$ cfu/µl (final concentration) of an overnight culture unless otherwise noted. The OD600 was measured every 10 minutes after infection for 24 hours using an automatic plate reader maintaining a temperature of 37°C. For curves of individual phage isolates (Fig. S2), each well was infected at an MOI of 1. For infections using communities 2, 3, 6 and 7 (Fig. S11), the phages were first passaged for 2 days to reach stable equilibrium. Growth curves were then measured after infecting bacteria with 5µl of phage communities which is the same dilution factor (1:30) used during passaging.

Culture age infection assay

To generate the 3-hour culture used for infection, a 24-hour culture was diluted 1:1000 in 20ml LB and grown for 3 hours to an OD of 0.1-0.3. This culture was then pelleted by centrifugation (10krpm for 3 minutes) and resuspended to an OD of ~5.0. The 72-hour culture was generated by inoculating a culture 3 days before the infection, while the 24-hour culture was started as previously described. We validated that the OD600 gave an accurate measurement for cfu concentration by serially diluting cultures onto LB plates and quantifying cfu. We were thus able to use the OD readings of each culture to infect the same number of cfu from each culturing condition. We then diluted each culture into fresh LB + salts ($4 \times 10^4$ cfu/µl final concentration) and infected with phage. For infections with a single phage, we used 10µl of each phage stock. For infections with community 2, we passaged phage N and S together for 2 days to reach equilibrium before infecting cells with 10µl. Phages were quantified after incubation through top agar plating.

Flow cytometer settings

The FACS Aria III cell sorter with a 100 µm nozzle was used for all flow cytometry experiments. Events were detected using forward scatter and side scatter triggers and data was obtained in logarithmic mode then analyzed with BD FACS software. The forward and side scatter settings (FSC at 6 volts, SSC at 300 volts) were chosen to avoid background noise from the buffer solution. Cultures were diluted 1:100 in PBS or LB + salt before sorting to ensure accurate cell separation. The PBS added to samples before sorting was passed through a 0.22 µm filter to reduce micro-particles in the buffer that could clog the machine or contribute to background noise.

Coinfected single-cell sorting

Cells were infected at various MOIs with phages from community 2 or 6 and incubated for 1.5 hours before sorting, which allowed time for phages to adsorb, but before the phages caused lysis (Fig. S11). Cells were gated to select for the entire bacterial population and then single-cell sorted into a 96-well PCR plate containing 20µl of LB + salt. Immediately after sorting the plate was sealed with pierceable aluminum foil and moved to 37C for 24h incubation. To determine the phages present, 5µl of each well was plated for the top agar assay.

A number of control experiments were performed to verify the suitability of this technique for determining the phage produced by individual cells. To verify that extracellular phages were not present in the drop together with a dispensed cell, we used unfiltered LB as a substitute for cells (this media contains particles with the same forward and side scatter as *E.* coli). We then added phage at the same concentrations as used in the infection experiments (see above). We sorted these background particles into a 96-well plate and checked for any plaques by plating using the top agar assay. Out of 120 wells we found 1 plaque in 3 different wells which confirmed that our error rate for counting free virus (instead of phage burst from a cell) was very low (1 plaque per 40 wells).

We then tested the accuracy of the machine for dispensing single bacterial cells by sorting an uninfected culture and plating the entire well on an LB plate. After overnight incubation of the agar plate, we found only 1 well out of 72 wells grew 2 colonies (none grew more than 2 colonies) (Fig. S8A). It is possible that these two colonies were produced not because of an error from sorting but instead due to cell division during the 30 minutes in between sorting and plating the sample. We also found that 19.4% of wells did not grow any colonies, suggesting that the flow cytometer had failed to sort a cell into those wells.

Size-based sorting and infection

To sort small and large cells, we gated the 24-hour culture's bacterial population by forward scatter capturing the smallest and largest 30% of cells. For growth curves of small and large cells, we sorted 100 cells from each size-gated population into 4 wells of a 96-well plate each containing 200µl LB. We then proceeded with the growth curve analysis as previously described. We confirmed the accuracy of our sorting by quantifying the cfu of each group immediately after sorting.

For assaying the production of phages in different subpopulations, we gated and sorted small and large populations using 2-way sorting at 10K events/second to sort a total population of 4 million cells from each gate into an empty 2ml collection tube. The total volume of sorted cells exceeded the 2ml maximum volume captured in the collection tube so we periodically emptied the collection tube into a 50ml falcon tube. Once sorting for both populations completed (~40 minutes) we reconcentrated the bacteria to ensure that phage infections would happen in a timely manner. To reconcentrate the bacteria we collected each sorted population onto a different 0.22 µm syringe disc filter fitted into a filter holder. Using sterile forceps we then moved the disc filter from the syringe holder into a 1.5ml tube containing 250µl LB + salts. The bacteria were released from the filter into the media by briefly vortexing. We then plated serial dilutions of the samples onto LB agar plates (before infection) to quantify cfu of the resuspended small and large populations. For both small and large populations we infected ~10k cfu with ~90k total pfu of community 2 in triplicate for a total reaction size of 25µl per well. After 24 hours of incubation we quantified the bacterial titer through cfu counting and filtered the infected samples. Bacterial abundances differed less than 1.3 fold between the small and large sized samples. We quantified the abundance of phage N and S produced by the small and large cell samples through top agar plating. Confidence intervals were calculated using standard error.

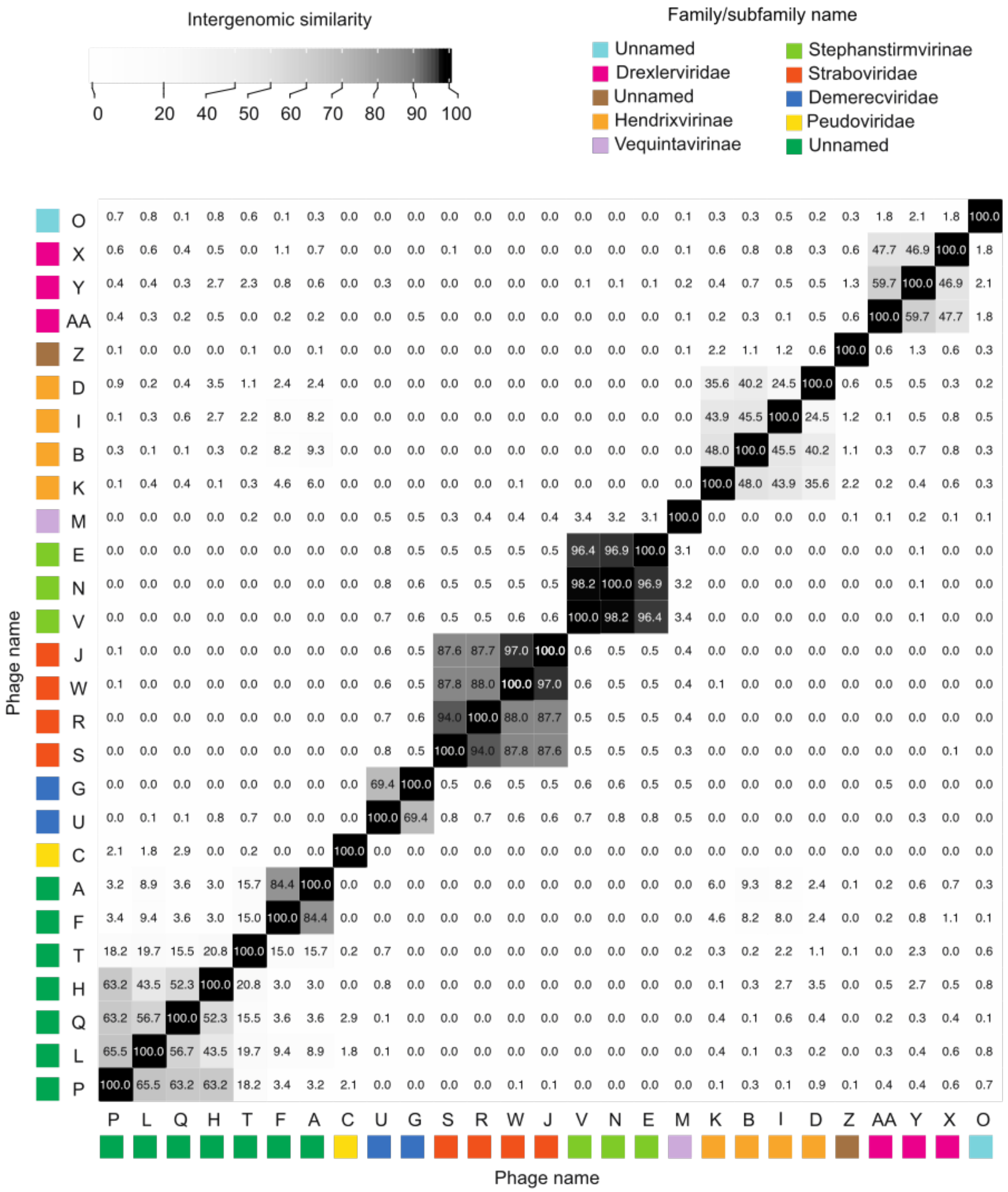


**Fig. S1. Phage genomes are unique, but have sequence homology within the same family.** Pairwise comparisons of all 27 genomes for the phages in our collection generated using VIRDIC (*58*). Darker colors indicate higher levels of intergenomic sequence similarity. Phage names are colored by their family classification, but not all families are named in ICTV (*28*), so subfamily names are listed in the legend in some cases. Phages were ordered along the x- and y-axis by sequence similarity, but this ordering naturally produced clustering among related phages in the same family. Phages within the same families are highly divergent from phages in other families (average <0.5% similarity in pairwise comparisons).

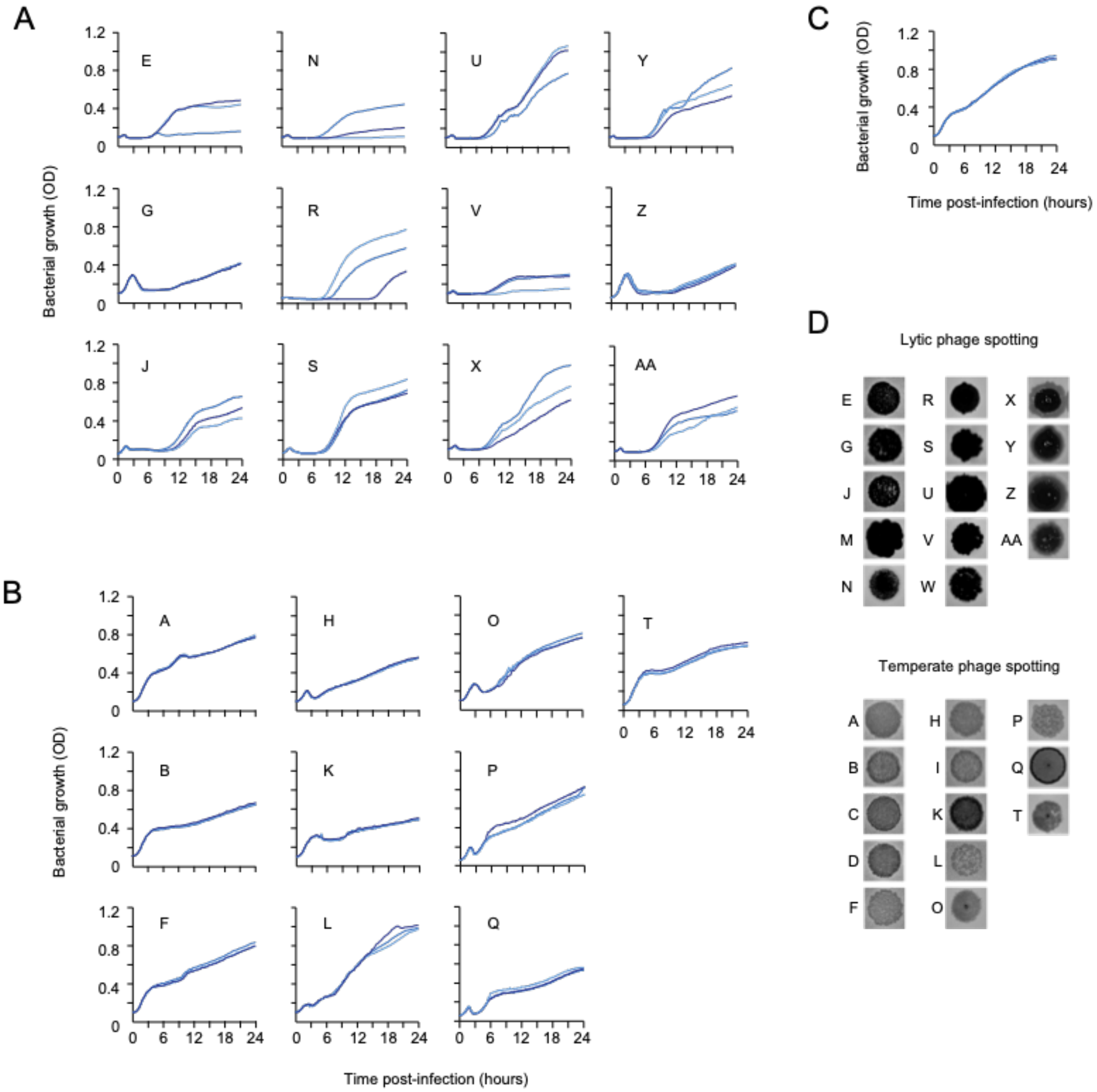


**Fig. S2. Growth curves and spot turbidity indicate the phage replication strategy.** (**A**) Bacterial density measured every 10 minutes for 24 hours after infection with a subset of lytic phages, with phage names inset within each growth curve. All phages reduce the OD to the limit of detection (OD ~0.1) within 3 hours post-infection and growth was remained undetectable for several hours, which was indicative that these were lytic phages. Regrowth 4 hours after lysis due to phage-resistant bacterial mutants that have emerged. (**B**) Same as (A) except a subset of temperate phages. Apart from phage Q, these phages did not reduce the density to the detectable limit at any point during infection. (**C**) Same as (A) except a no phage control (**D**) High titer phage stocks were spotted on a lawn of *E. coli* to characterize their replication strategy, since temperate phages tend to produce more turbid spots. These classifications were also confirmed bioinformatically (Methods).

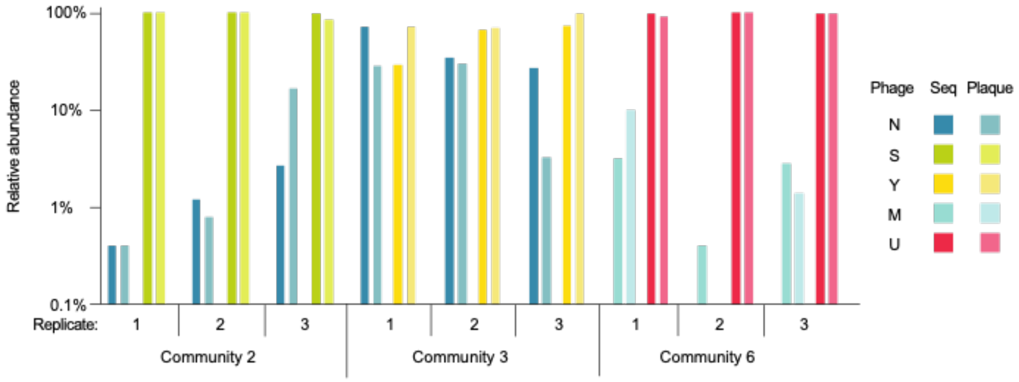


**Fig. S3. The abundance and identity of communities was similar when determined through plaque counting and deep sequencing.**

The replicates for three communities were plated on top agar at the final passage to determine the relative abundance of each phage through plaque morphology (Plaque) and then compared to the relative abundances calculated through deep sequencing (Seq).

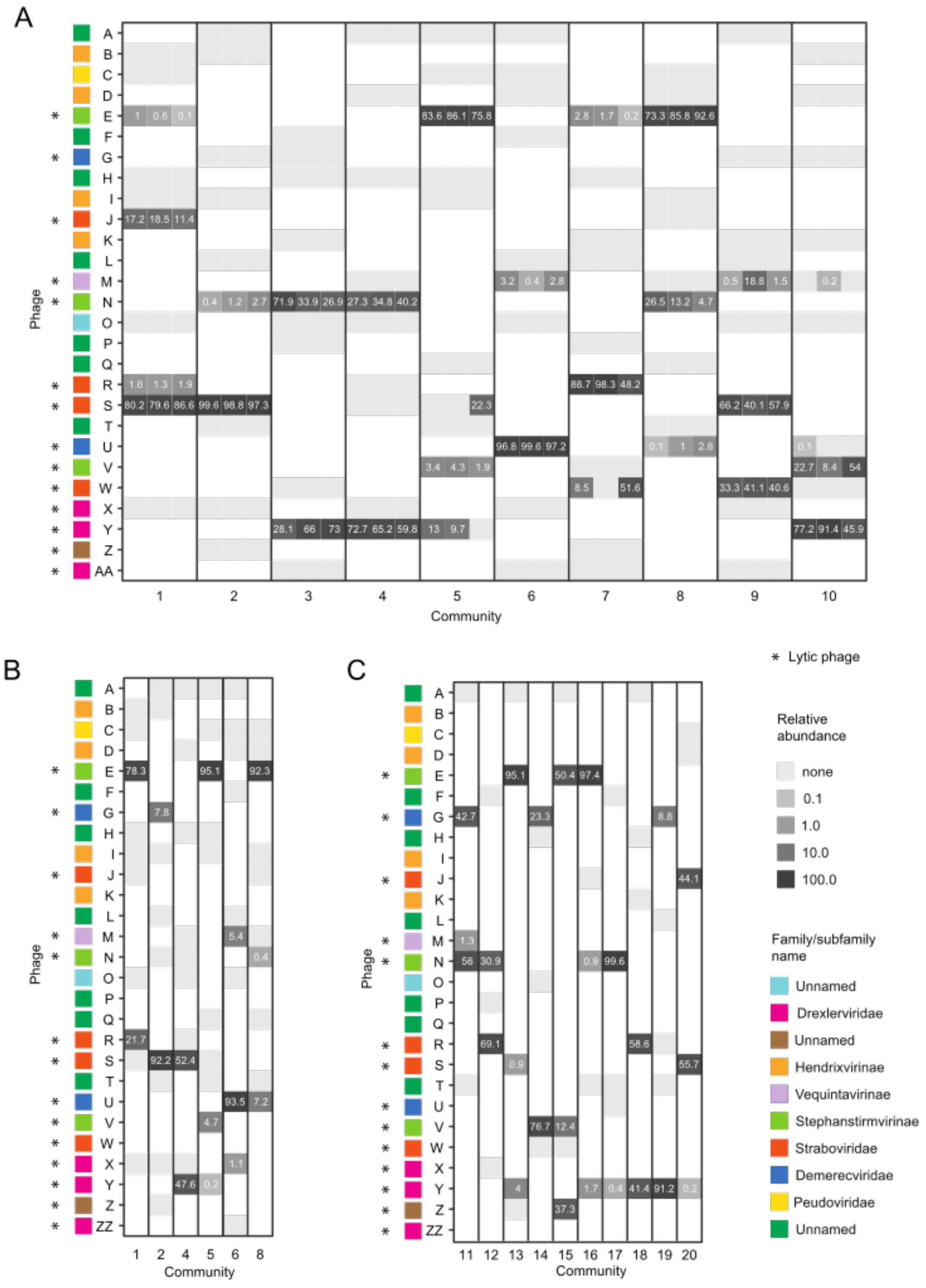
A
B
C
Phage
Community
* Lytic phage
Relative abundance
none
0.1
1.0
10.0
100.0
Family/subfamily name
Unnamed
Drexlerviridae
Unnamed
Hendrixvirinae
Vequintavirinae
Stephanstirmvirinae
Straboviridae
Demerecviridae
Peudoviridae
Unnamed

**Fig. S4. Relative abundances for phages found in all communities and replicates.**
(**A**) The relative abundance of each phage was determined through deep sequencing for all communities and replicates. The relative abundance of each phage was calculated based on the average coverage of each phage's genome. Darker colors indicate higher abundance, with color scaled by log relative abundance. Phages with "none" were added in the initial community but not detected through deep sequencing at the final passage. The limit of detection was ~0.1% of the total population. (**B**) Same as (A) except communities were passaged without filtering E. coli between each passage and only a single replicate was performed for each community. The starting communities used in this experiment were composed of the same phage strains as communities 1, 2, 4-6, and 8 used in (A). These final communities all had similar compositions (with some notable differences) as communities passaged with filtering. (**B**) Same as (A) except the initial communities contained 5 instead of 10 phages and only a single replicate was performed for each community.

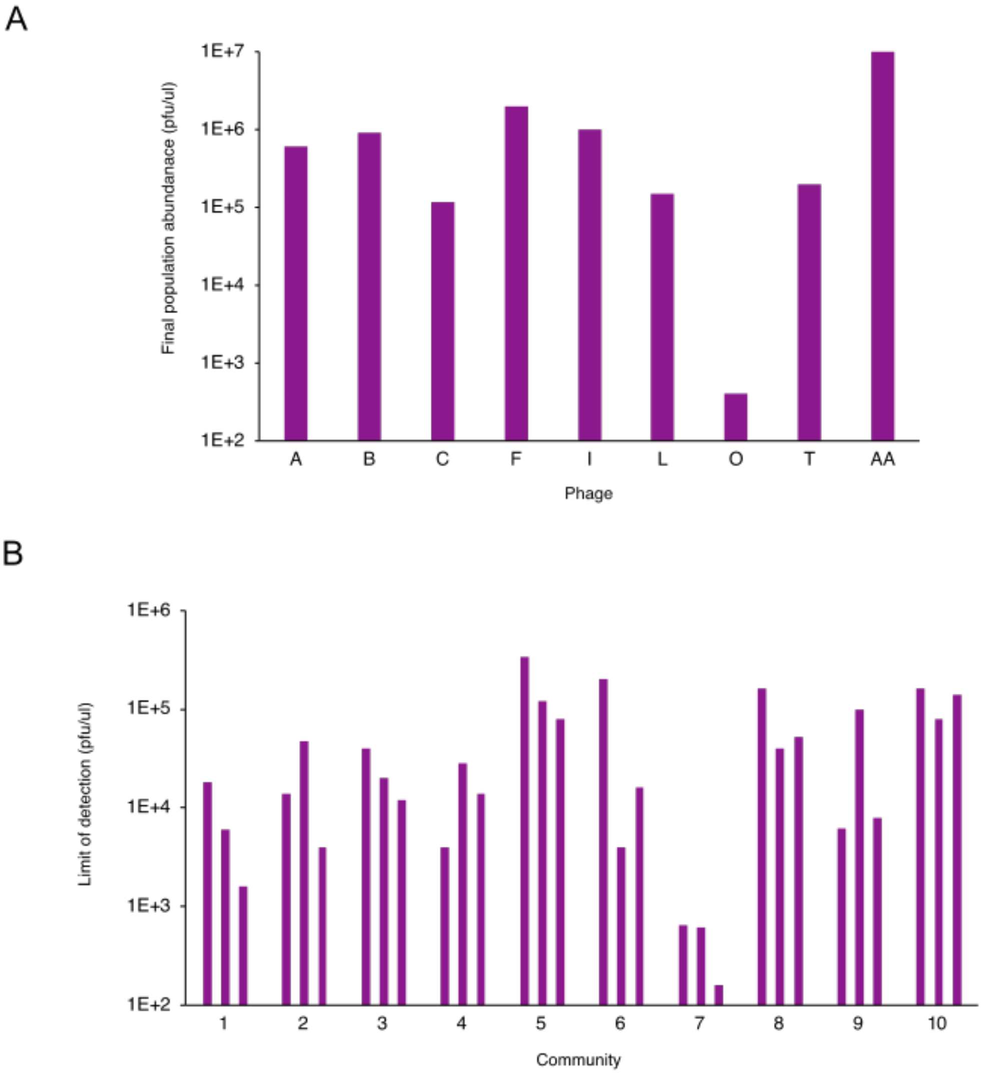


**Fig. S5. Temperate phages survive passaging in isolation.**
(**A**) Final abundances of a subset of temperate phages, and lytic phage AA which was absent from any final community, when passaged every 24 hour for 12 days with filtering. All phages were detectable at the last passage. (**B**) The limit of detection for phages in our community was calculated as around 0.1% of the final abundance of phages in the community, as determined through top agar plating, because the limit of detection through deep sequencing was around 0.1% of the total abundance of phages in our communities. In most cases, this limit of detection was greater than the phage titers during passaging in (A), suggesting that if these phages had persisted in community they would have been detectable through sequencing.

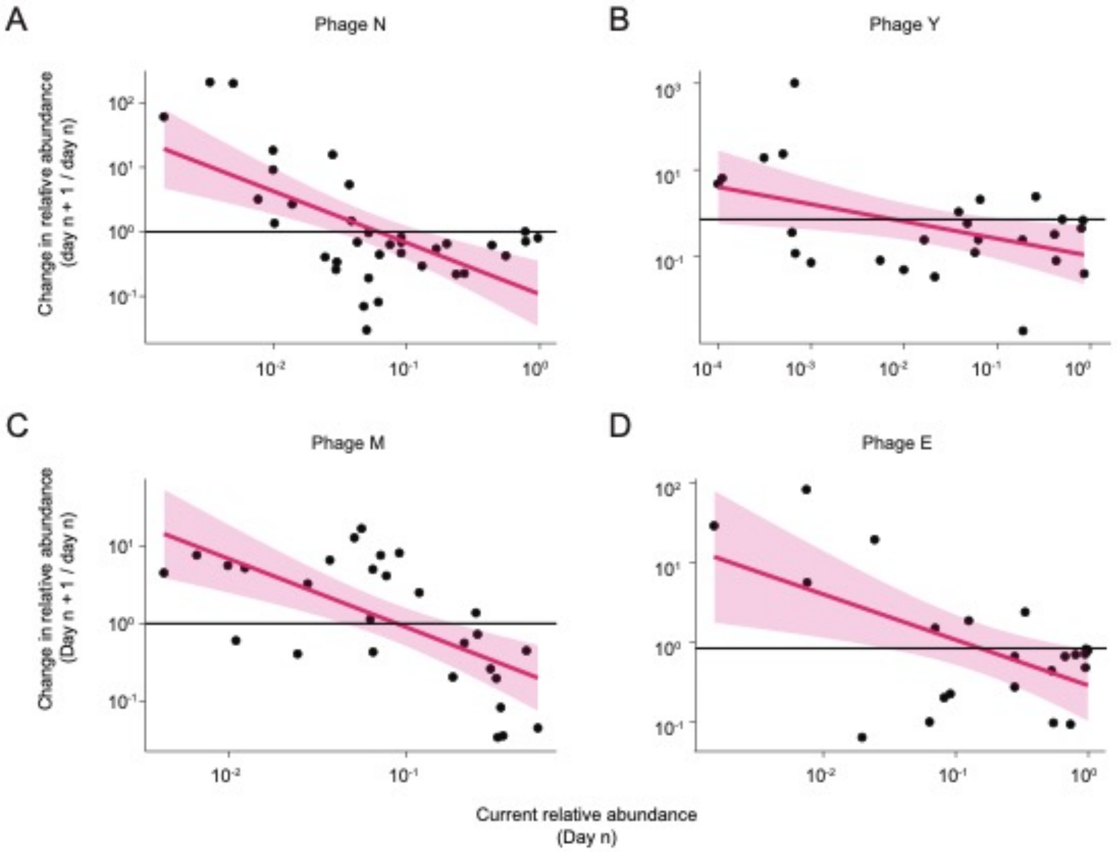


**Fig. S6. Phages in community are under negative frequency dependent selection.**
The change in relative abundance of one of the phages in a community (listed in the subtitle) is plotted against that phage's change in relative abundance using data from Fig. 2B for (**A-D**) communities 2, 3, 6, and 7. The y-axis is calculated by dividing the relative abundance of a phage on the following passage (n + 1) by the abundance of that phage at the current passage (n). For all four communities, this relationship was negative, indicating that both phages in the population did better when rare (negative frequency dependence). In addition, for all four communities, the line crosses zero, indicating that a stable equilibrium containing both phages exists. The solid pink line is was fitted to the data with a linear regression model and the shaded pink area indicates the 95% confidence interval. The solid black line crosses the y-axis at 1, indicating where no change in proportion is predicted, ie. when the future proportion and current proportion are equal to 1.

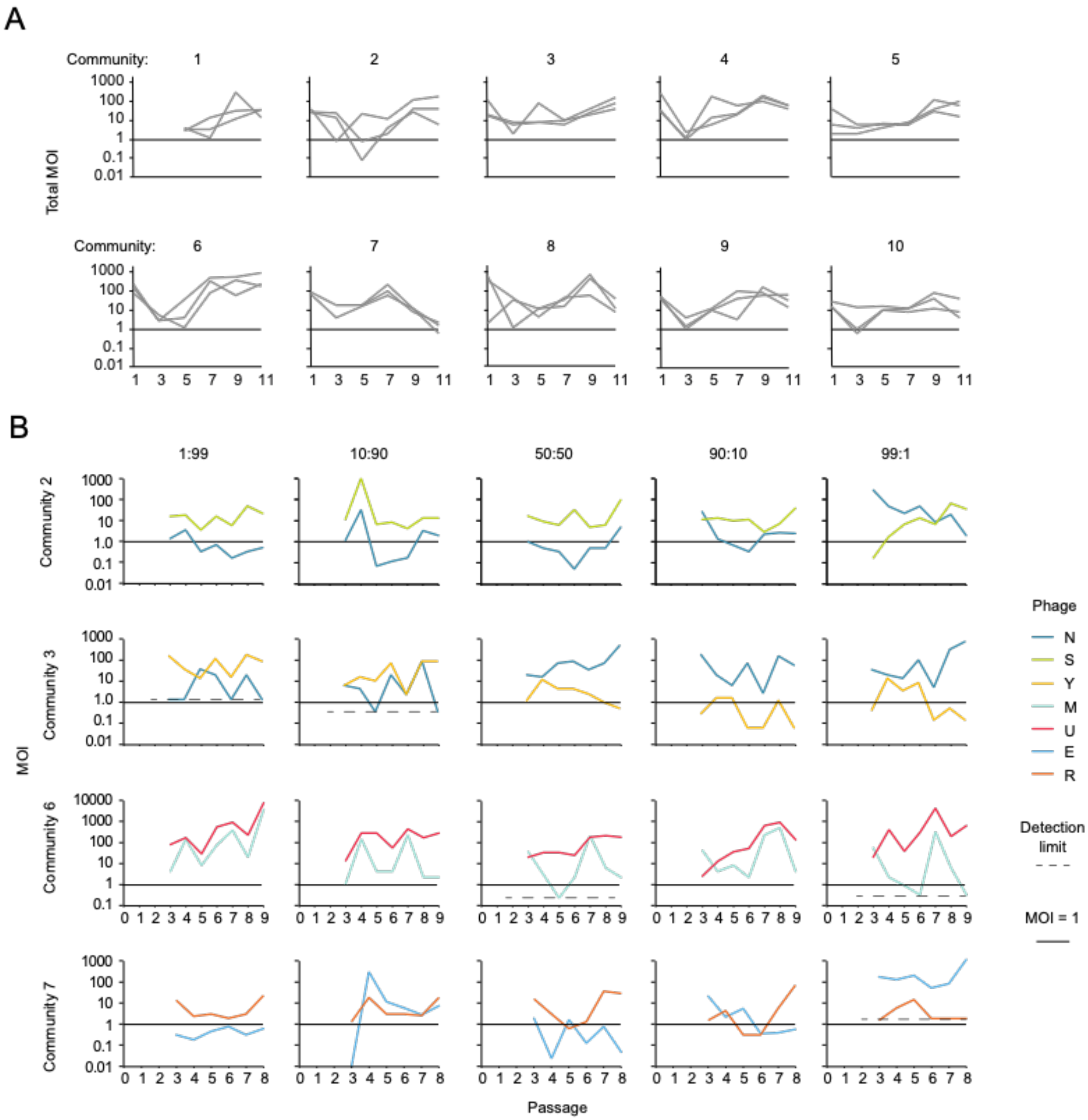


**Fig. S7. Phages are more abundant than bacteria at the onset of each passage.**
The ratio of phage to bacteria (MOI) for each community at the onset of each passage. The abundance of bacteria was calculated based on the average density of an overnight culture (OD of 5). The abundance of phage was calculated through phage titering for (**A**) three replicates of all 10 communities in Fig. 1e and (**B**) samples in Fig. 2B.

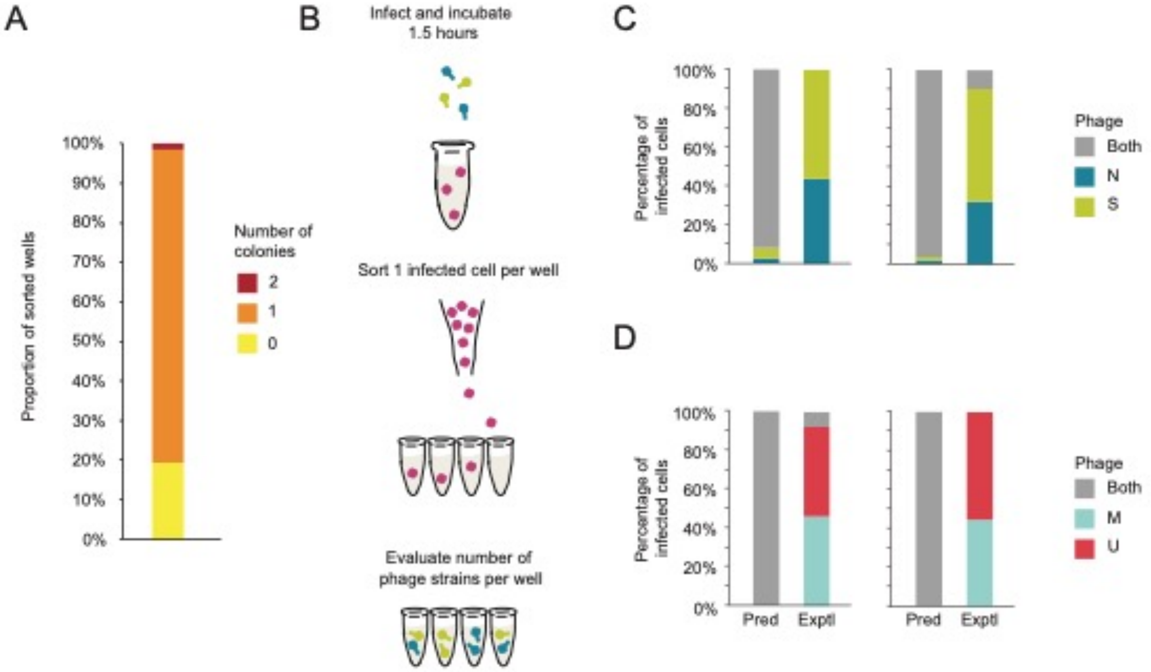


**Fig. S8. Multiple phage species rarely reproduce within the same cells.**
(**A**) Validation that flow cytometry accurately sorts individual bacterial cells. The proportion of wells that produced 0, 1, or 2 colonies after flow cytometry sorting. A single event was sorted into a well containing 20ul of LB and then the entire contents of the well was plated onto LB agar and incubated to determine colony formation. A total of 72 wells were counted.
(**B**) Schematic for determining the phages species produced from a single coinfected cell. Co-infected cells were sorted into wells containing media just before lysis. Each well was then plated and the phage species produced by each coinfected cell was determined through top agar plating. (**C**) Successful coinfection is rare, as seen in the prevalence of coinfected cells that produce one or two phage strains. The predicted (pred) values were calculated using a Poisson distribution based on the MOI used during the experiments. The experimental results (exptl) were collected at 2 different MOIs for community 2 (left: MOI of 3.9, n = 16 cells; right: MOI of 4.8, n = 31 cells) and (**D**) community 7 (left: MOI of 25, n = 24 cells; right, MOI of 14, n = 9 cells).

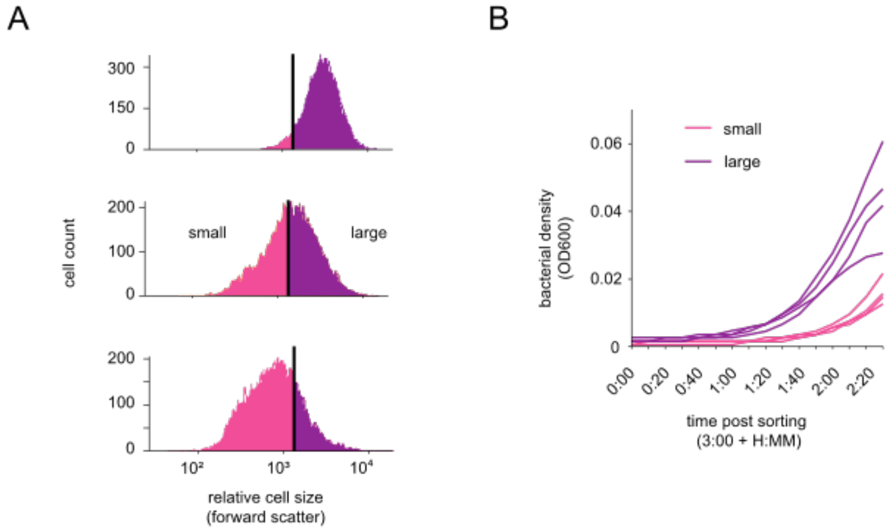


**Fig. S9. Cell size is an accurate marker for growth physiology.**
(**A**) Histograms of relative cell size for 3-hour, 24-hour, and 72-hour cultures (top to bottom), as determined by flow cytometry, showing that cell size decreases with increasing culture age (**B**) Growth curves of small or large cells from a 24-hour culture show growth differences. Small or large cells (100 cells per replicate) were sorted into 4 wells of a 96-well plate containing LB media by gating events with the highest 30% (large cells) or lowest 30% (small cells) forward scatter. OD was measured every 10 minutes for 24 hours using an automated plate reader.

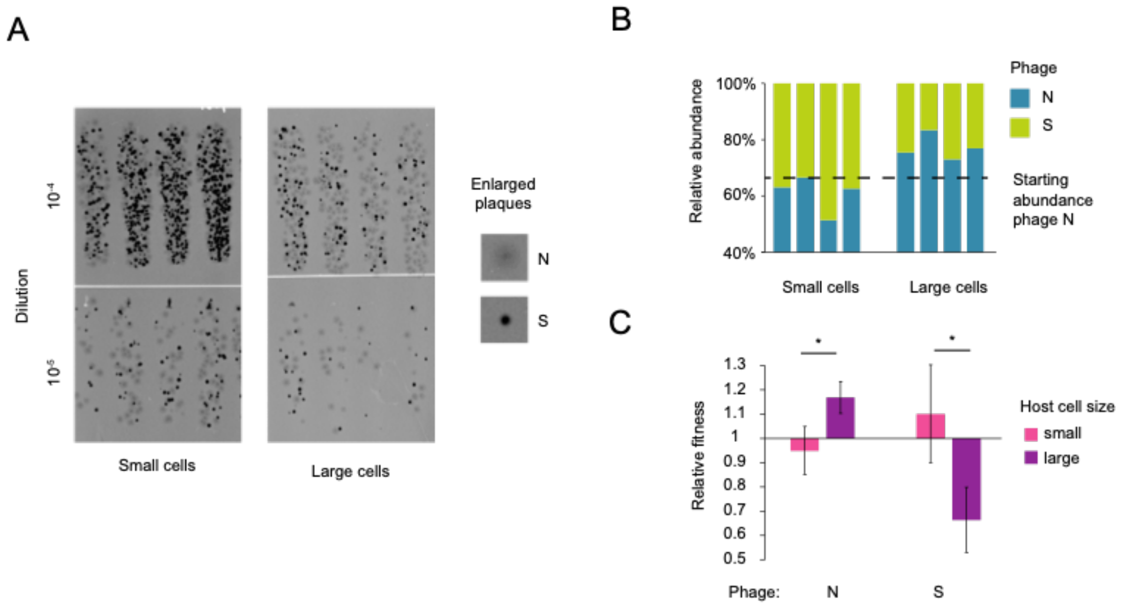


**Fig. S10. Small and large bacterial subpopulations preferrentially produce different phage strains.**

Replicate experiment of Fig. 4D. (**A**) Top agar plating of phages produced by cells of different sizes (**B**) Relative abundance of each phage, with the dotted line indicating the starting proportion of phage N (**C**) The population change of each phage by cell size, relative to the starting proportion. Results are the mean of four biological replicates $P$ = .006 from a two-tailed T test.

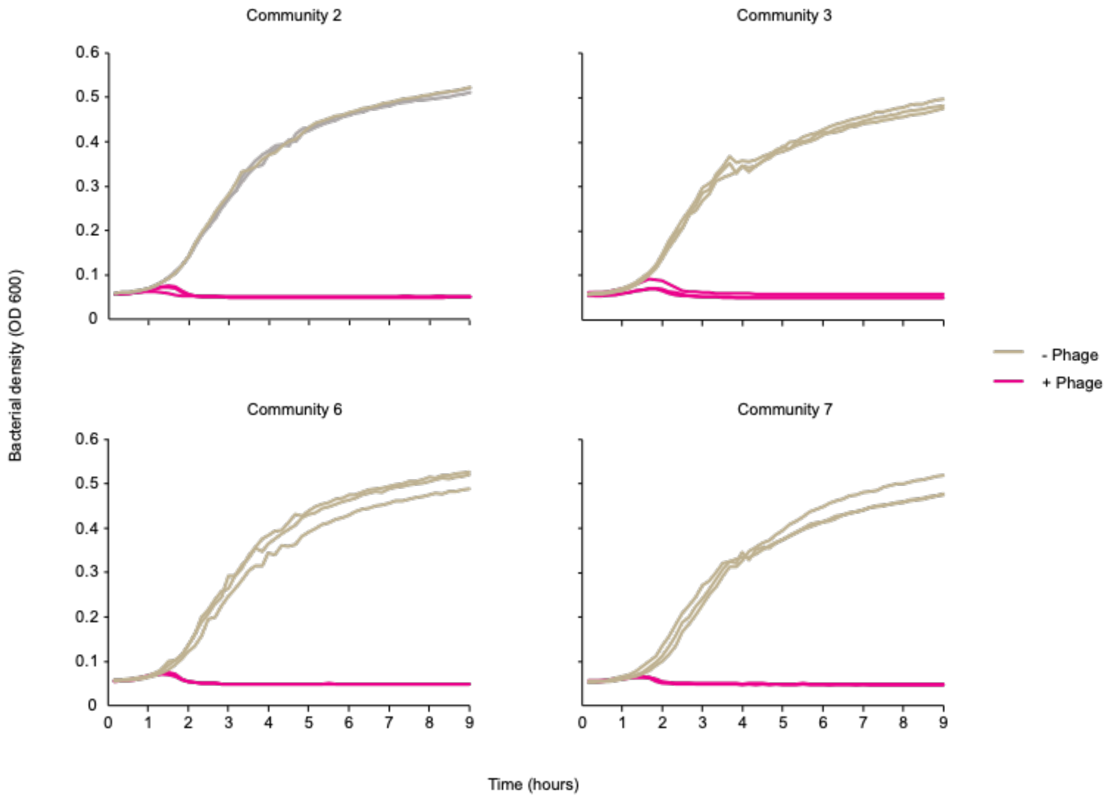


**Fig. S11. Bacterial lysis occurs shortly after the onset of infection.**
Bacterial densities measured every 10 minutes during 9 hours after infection for triplicate wells. For samples with phage, the initial increase in OD was followed by lysis around 1-2 hours after infection. No subsequent increase in density was observed throughout the 24 hour incubation (even past the 9 hours shown here).

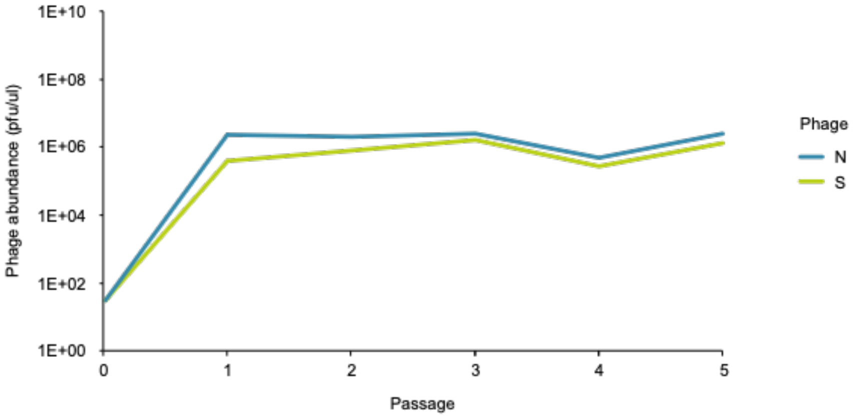


**Fig. S12. Changing the bacterial culture inoculum increased the stability of community 2.** Passaging of community 2 on an overnight culture started from a frozen stock gave greater stability than when this community was passaged on a culture started from a single colony (compare to community 2 in Fig. 2a, b). Abundances of each phage were determined through top agar plaquing.

**Table 1. Phage collection characteristics.** Lytic and temperate bioinformatics classifications refer to (A) BACPHLIP (*60*) and (B) PhaTYP (*59*).

| | | | Taxonomic Classification | | | | Lytic | | Temperate | |
|---|---|---|---|---|---|---|---|---|---|---|
| Short name | Pres/Abs from communities | Replication | ICTV family | ICTV subfamily | ICTV genus | Formal name | A | B | A | B |
| A | absent | temperate | Unnamed | Unnamed | Unnamed | Escherichia phage Eeyore | 0.05 | | 0.95 | 0.9985 |
| B | absent | temperate | Unnamed | Hendrixvirinae | Unnamed | Escherichia phage LMessi | 0.00 | | 1.00 | 0.9999 |
| C | absent | temperate | Peduoviridae | Unnamed | Unnamed | Escherichia phage Chuche | 0.00 | | 1.00 | 0.9998 |
| D | absent | temperate | Unnamed | Hendrixvirinae | Unnamed | Escherichia phage SaltyDalty | 0.00 | | 1.00 | 0.9999 |
| E | present | lytic | Unnamed | Stephanstirmvirinae | Justusliebigvirus | Escherichia phage Antha | 0.44 | 0.9992 | 0.56 | |
| F | absent | temperate | Unnamed | Unnamed | Unnamed | Escherichia phage Flotzy | 0.05 | | 0.95 | 0.9998 |
| G | present | lytic | Demerecviridae | Markadamsvirinae | Epseptimavirus | Escherichia phage SpoonbillGilly | 0.96 | 0.9999 | 0.04 | |
| H | absent | temperate | Unnamed | Unnamed | Unnamed | Escherichia phage IdasGreatGrand | 0.00 | | 1.00 | 0.9990 |
| I | absent | temperate | Unnamed | Hendrixvirinae | Unnamed | Escherichia phage CharmingMan | 0.00 | | 1.00 | 0.9998 |
| J | present | lytic | Straboviridae | Tevenvirinae | Tequatrovirus | Escherichia phage Pinarejo | 1.00 | 0.9999 | 0.00 | |
| K | absent | temperate | Unnamed | Hendrixvirinae | Unnamed | Escherichia phage Fostie | 0.10 | | 0.90 | 0.9998 |
| L | absent | temperate | Unnamed | Unnamed | Unnamed | Escherichia phage Kati | 0.00 | | 1.00 | 0.9991 |
| M | present | lytic | Unnamed | Vequintavirinae | Vequintavirus | Escherichia phage MiataMamis | 0.90 | 0.9999 | 0.10 | |
| N | present | lytic | Unnamed | Stephanstirmvirinae | Justusliebigvirus | Escherichia phage Nernie | 0.45 | 0.9998 | 0.55 | |
| O | absent | temperate | Unnamed | Unnamed | Unnamed | Escherichia phage MagusPhagus | 0.05 | | 0.95 | 0.9998 |
| P | absent | temperate | Unnamed | Unnamed | Unnamed | Escherichia phage Turner | 0.00 | | 1.00 | 0.9993 |
| Q | absent | temperate | Unnamed | Unnamed | Unnamed | Escherichia phage Chlobot | 0.00 | | 1.00 | 0.9995 |
| R | present | lytic | Straboviridae | Tevenvirinae | Tequatrovirus | Escherichia phage PinkRuby | 0.86 | 0.9999 | 0.14 | |
| S | present | lytic | Straboviridae | Tevenvirinae | Tequatrovirus | Escherichia phage SholemSchwarzbard | 0.91 | 0.9999 | 0.09 | |
| T | absent | temperate | Unnamed | Unnamed | Glaedevirus | Escherichia phage HannyBb | 0.00 | | 1.00 | 0.9992 |
| U | present | lytic | Demerecviridae | Markadamsvirinae | Tequintavirus | Escherichia phage KookSpook | 0.96 | 0.9999 | 0.04 | |
| V | present | lytic | Unnamed | Stephanstirmvirinae | Justusliebigvirus | Escherichia phage VivaForever | 0.44 | 0.9998 | 0.56 | |
| W | present | lytic | Straboviridae | Tevenvirinae | Tequatrovirus | Escherichia phage Elenatom | 1.00 | 0.9999 | 0.00 | |
| X | present | lytic | Drexlerviridae | Tempevirinae | Unnamed | Escherichia phage BananaBelafonté | 0.84 | 0.9999 | 0.16 | |
| Y | present | lytic | Drexlerviridae | Tempevirinae | Warwickvirus | Escherichia phage BabyEgg | 0.99 | 0.9999 | 0.01 | |
| Z | present | lytic | Unnamed | Unnamed | Dhillonvirus | Escherichia phage Zumzum | 0.95 | 0.9999 | 0.05 | |
| AA | absent | lytic | Drexlerviridae | Tempevirinae | Warwickvirus | Escherichia phage Kalemo | 1.00 | 0.9999 | 0.00 | |
